# Detection of threshold-level stimuli modulated by temporal predictions of the cerebellum

**DOI:** 10.1101/2023.06.20.545246

**Authors:** Lau M. Andersen, Sarang Dalal

**Affiliations:** Center of Functionally Integrative Neuroscience (CFIN), Aarhus University, Universitetsbyen 3, Building 1710, 8000 Aarhus C, Denmark; Aarhus Institute of Advanced Studies (AIAS), Aarhus University, Høegh-Guldbergs Gade 6B, 8000 Aarhus C, Denmark; Department for Linguistics, Cognitive Science and Semiotics, Aarhus University, Jens Chr. Skous Vej 4, Building 1485, 8000 Aarhus C, Denmark

## Abstract

The cerebellum has the reputation of being a primitive part of the brain that mostly is involved in motor coordination and motor control. Older lesion studies and more recent electrophysiological studies have, however, indicated that it is involved in temporal perception and temporal expectation-building. An outstanding question is whether this temporal expectation-building cerebellar activity has *functional relevance.* In this study, we collected magnetoencephalographic data from 30 healthy participants performing a detection task on at-threshold stimulation that was presented at the end of a sequence of temporally regular or irregular above-threshold stimulation.

We found that behavioural detection rates depended on the degree of irregularity in the sequence preceding it. We also found cerebellar responses evoked by above-threshold and at-threshold stimulation. The evoked responses to at-threshold stimulation differed significantly, depending on whether it was preceded by a regular or an irregular sequence. Finally, we found that detection performance across participants correlated significantly with the differences in cerebellar evoked responses to the at-threshold stimulation, demonstrating the functional relevance of cerebellar activity in sensory expectation-building.

We furthermore found evidence of thalamic involvement, as indicated by responses in the beta band (14-30 Hz) and by significant modulations of cerebello-thalamic connectivity by the regularity of the sequence and the kind of stimulation terminating the sequence.

These results provide evidence that the temporal expectation-building mechanism of the cerebellum, what we and others have called an internal clock, shows functional relevance by regulating behaviour and performance in sensory action that requires acting and integrating evidence over precise time-scales.

## 1 Introduction

The cerebellum is being studied with renewed interest after it has become clear that its function is not only one of motor coordination, but also activity related to cognition (Andersen et al., 2020; King et al., 2019). Part of its cognitive abilities have been thought to be that of detecting stimulus rhythmicity (Ivry, 1997). In humans, evidence for this ability has come from lesion studies of cerebellar patients (Ivry, 1993). Recent studies, however, have provided electrophysiological, non-invasive evidence of cerebellar involvement in temporal sensory prediction (Andersen and Dalal, 2021; Andersen and Lundqvist, 2019; Herrojo Ruiz et al., 2017). The trust in these findings is warranted by a recent study demonstrating that both electroencephalography (EEG) and magnetoencephalography (MEG) are sensitive to cerebellar activations (Samuelsson et al., 2020). These studies have, however, only shown evidence for cerebellar activity correlating with temporal predictability, not for the functional relevance of cerebellar activity in potentially guiding action. To show functional relevance, it is necessary to show that it influences behaviour. Given such an influence on behaviour, it is natural to ask what neural connections would make it possible to elicit informed motor action. Connections from the cerebellum to the primary motor cortex would indicate that such an influence is possible. Such a connection has been reported in monkeys to take place through the thalamus (Bostan et al., 2013; Orioli and Strick, 1989). Furthermore, evidence of functional connectivity between the cerebellum, thalamus and the primary motor cortex has been reported in natural tremor and essential tremor using magnetoencephalography (Gross et al., 2002; Pollok et al., 2008; Schnitzler et al., 2009). Thus, it is possible that the thalamus is instrumental in making sensory expectations in the cerebellum informative to behaviour due to its connections through the thalamus to the primary motor cortex. Besides anatomical connections to the thalamus, anatomical connections have also been reported between the cerebellum, the thalamus and the basal ganglia (Bostan et al., 2013; Bostan and Strick, 2010). This is interesting because basal ganglia are thought to be involved in decision making (Bogacz and Gurney, 2007; Ding and Gold, 2013). Interestingly, we earlier found evidence of thalamic responses (Andersen and Dalal, 2022: Supplementary Material 3: New Figure 3) in the beta band (14-30 Hz). This makes it possible to test whether the cerebellum and the thalamus show functional connectivity, if we replicate the finding of thalamic activity in the beta band.

The primary hypothesis of this study is to show that cerebellar activity reflective of temporal sensory expectations is informative to behaviour. An exploratory sub-hypothesis is that the thalamus and will be connected to cerebellum at the functional level.

To operationalise these hypotheses, we needed a paradigm contrasting temporal expectations that would both elicit differences in behavioural performance and in cerebellar activity. We adapted the passive paradigm of Andersen and Dalal (2021), with which we showed that cerebellar activity in the beta band (14-30 Hz) elicited by unexpectedly omitted stimulations correlates with the precision of temporal expectations. In making the paradigm active, we had participants decide whether or not at-threshold electrical stimulation was applied to their finger or whether stimulation was omitted following a train of either precisely presented suprathreshold stimulation or a train of temporally jittered suprathreshold stimulation.

We thus expect for the primary hypothesis that: 1) the finding that cerebellar activity in the beta band (14-30 Hz) for unexpectedly omitted stimulations correlating with the precision of temporal expectations will replicate; 2) participants will detect at-threshold stimulation at a higher rate when these stimulations are preceded by trains of precisely presented suprathreshold stimulation as compared to when they are preceded by trains of temporally jittered suprathreshold stimulation; and 3) participants’ detection performance will correlate with cerebellar activity evoked by the at-threshold stimulation.

In terms of the sub-hypotheses, we expect that the cerebellum will show differential patterns of functional connectivity to the thalamus for the at-threshold stimulation and omissions at the end of trains of stimulation, depending on whether these are precisely presented or temporally jittered.

## 2 Materials and Methods

### 2.1 Participants

Thirty right-handed, healthy participants volunteered to take part in the experiment (twenty-two male and eight female, Mean age: 29.6 y; Standard Deviation: 6.3 y; Range 18-43 y). The experiment was advertised in a local database that participants previously registered to. The experiment was approved by the local institutional review board in accordance with the Declaration of Helsinki. The participants provided informed consent and were compensated with 400 DKK for their participation. Two participants were excluded, one due to aborting the experiment due to discomfort and one due to excessive noise that led to rejection of more than 66% of the trials.

### 2.2 Stimuli and procedure

Tactile stimulation was generated by two ring electrodes driven by an electric current generator (Stimulus Current Generator, DeMeTec GmbH, Germany) The ring electrodes were fastened to the tip of the right index finger. One was placed 10 mm below the bottom of the nail and the other one 10 mm below that. Stimulation was applied in sequences of six stimuli, with an inter-stimulus interval of 1,497 ms, meaning that the stimulation would not lock to the 50 Hz power from the electrical environment. All pulses were 100 µs in duration. The current applied was individualised based on a staircasing procedure. The sixth stimulus was followed either by an omission of stimulation or a weak stimulation, the target stimulation. After the seventh stimulation, omitted or weak, participants were to make a response, indicating with a button press on a response box held in their left hand, as to whether or not they were stimulated, allowing a categorisation of their responses into *hits, misses, false alarms* or *correct rejections* (MacMillan, 2002). The next sequence would begin 1,497 ms after their button press.

The six stimuli leading up to the target stimulation were either presented in a no-jitter sequence or a jittered sequence. In the no-jitter sequence, all six stimuli were exactly 1,497 ms apart (0% jitter). In the jittered sequence, the fourth, fifth and sixth stimuli had a jitter of up to 15% applied to them, i.e. they happened from -225 ms to 225 ms (in steps of 1 ms) relative to when the stimulus would have occurred, had the sequence been non-jittered. For each jittered stimulation, the jitter, an integer number of milliseconds, was chosen randomly from a uniform distribution. The number of sequences ending in either weak or omitted stimulations was counterbalanced such that an equal number of sequences was applied for each of the four possible sequences, (1. non-jittered ending with an omission, 2. non-jittered ending with a weak stimulation, 3. jittered ending with an omission, 4. jittered ending with a weak stimulation) (Figure 1).

**Figure 1:**
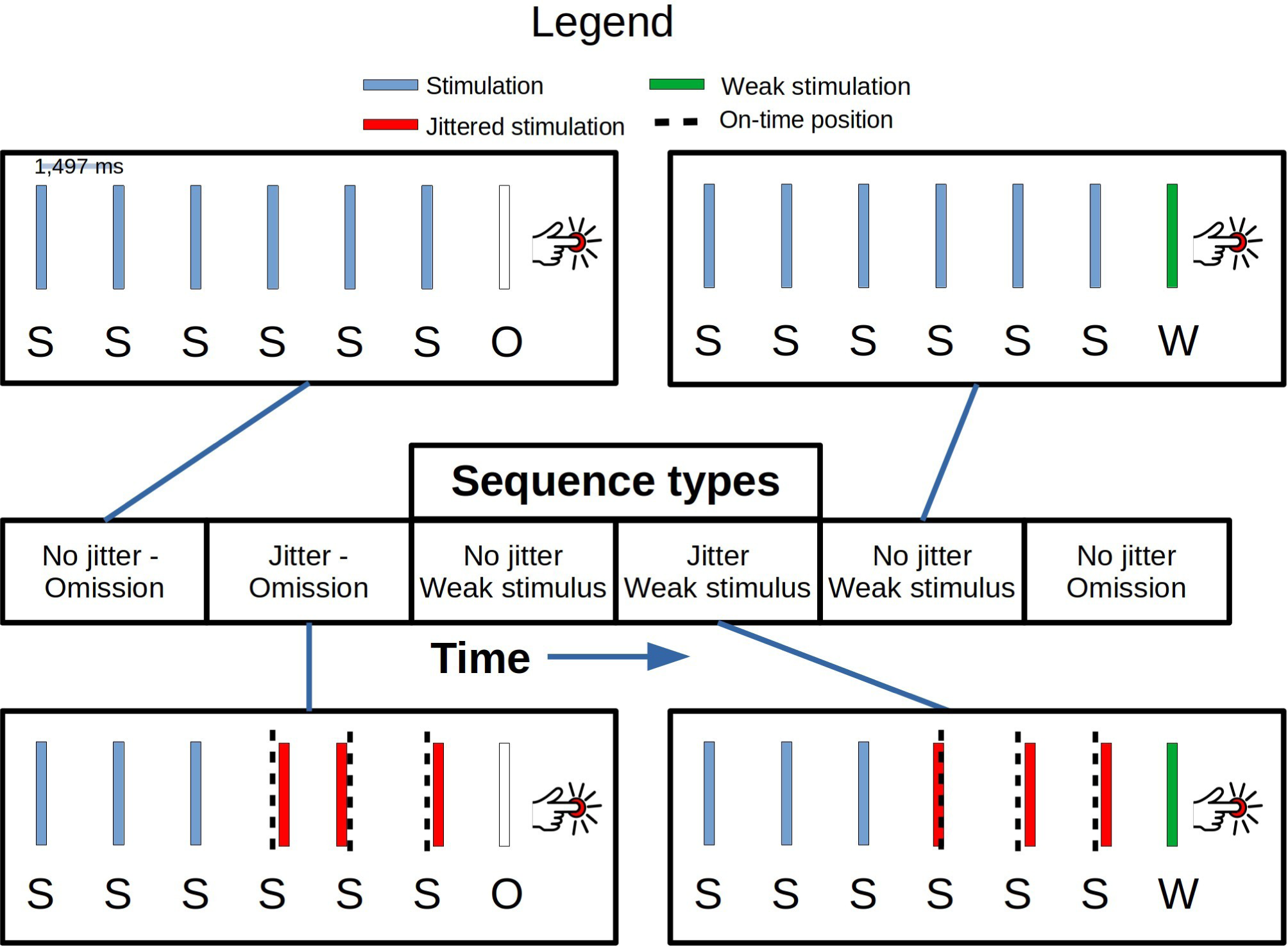
The paradigm: Four kinds of sequences were administered in a pseudorandom and counterbalanced order. The first three stimulations (S) were always on time, spaced 1,497 ms apart. In the jittered sequence, up to 225 ms of jitter in either direction (forwards or backwards in time) was applied to the following three stimulations. The red S-bars indicate jittered stimulations and the dashed black vertical lines indicate when the stimulation should have occurred, had the sequence been regular. At the seventh position, either an omission (O) or a weak (W) at-threshold stimulation would occur. Participants then had to indicate with a button press whether or not they had been stimulated at the time of the seventh position.

300 stimulation sequences were administered, 75 of each type presented in a pseudorandom order. This structure resulted in 300 *First Stimulations*, 300 *Second Stimulations* and 300 *Third Stimulations*, due to the first three stimulations being identical between the sequences. Furthermore, this resulted in 2 x 150 *Fourth Stimulations*, *Fifth Stimulations* and *Sixth Stimulations*, 150 for the non-jittered sequence and 150 for the jittered sequence. This structure finally gave to rise 2 x 75 *Omitted Stimulations* (*Omission 0* and *Omission 15)* and 2 x 75 *Weak Stimulations* (*Weak 0* and *Weak 15*), 75 of each for the non-jittered sequence and 75 each for the jittered sequence. The experiment consisted of 12 blocks – between each block, verbal contact was initiated to ensure that the participant was comfortable. PsychoPy3 (version 2020.2.3) (Peirce et al., 2019) was used to deliver the stimuli from a Linux operating system running Ubuntu Mate (kernel version: 4.15.0-154-generic). Electro-oculography, -cardiography and -myography were recorded to detect eye blinks, eye movements, the heart beat and muscle activity over the splenius muscles. For explorative purposes, respiration was measured using a respiratory belt for the last twenty-two participants.

### 2.3 Preparation of participants

In preparation for the measurement, each participant had two pairs of electrodes placed horizontally and vertically around the eyes. The heart beat was measured by having a pair of electrodes on each collarbone. Two pairs of electrodes were placed on either side of the splenius muscles. Four head-position indicator coils, two behind the ears and two on the forehead, were placed on the participants. The ground electrode was placed on the right wrist. We subsequently digitised the head shape of each participant using a Polhemus FASTRAK (Colchester, Vermont, USA). Three fiducial points, the nasion and the left and right pre-auricular points were alongside the positions of the head-indicator coils. Furthermore, we acquired around 150 extra points digitising the head shape of each participant. Participants were subsequently placed in the supine position of the magnetoencephalographic system and we took great care in making sure that they would lie comfortably, thus preventing neck tension. The ring electrodes were then put on the participant’s finger and the respiratory belt around their abdomen. Before beginning the actual experiment two sessions were run, a staircasing session and a detection session. The staircasing session was a 2-alternative forced choice session, where first the number *1* was shown on a screen and the the number *2* was shown, 1,497 ms between them. Either the number *1* or the number *2* was accompanied by a current. The participant then had to indicate with a button press when they were stimulated, either at the first or second position. 40 stimuli were applied. Each time they answered correctly, the current got less intense, and vice versa more intense each time they answered wrongly. A response prompt appeared on screen reminding them to answer. A psychometric curve was estimated from the 40 trials and the current estimated to be equivalent to answering correct on 85 % of the trials were chosen as the value for the weak stimulation in the actual experiment.

Subsequently, a detection session was run with 50 trials where they were stimulated with the individualised, staircased current on around half the trials and where no stimulation occurred on the other trials. Using the response box, they had to indicate whether or not they perceived a stimulus. If a participant ended up having more misses than hits in this session, the staircase and the detection sessions were run again before going on the actual experiment. A response prompt appeared reminding them to answer.

In the actual experiment, participants were instructed to fixate on a shown fixation cross and not move their bodies or heads. No response prompt occurred during the actual experiment (to prevent visually evoked responses). Participants were therefore instructed to count the first six stimuli internally before providing the appropriate response, as to whether a weak or omitted stimulation was presented.

### 2.4 Acquisition of data

Magnetoencephalographic data was recorded on an Elekta Neuromag TRIUX system inside a magnetically shielded room (Vacuumschmelze Ak3b) at a sampling frequency of 1,000 Hz. Data was acquired with low-pass and high-pass filters applied, at 330 Hz and 0.1 Hz respectively.

### 2.5 Processing of magnetoencephalographic data

For each participant, we inspected the raw data traces and their power spectra densities for bad sensors. Sensors were removed if they showed a consistently higher or lower power than the average. We analysed both evoked and oscillatory responses.

All magnetoencephalographic analyses were carried out in MNE-Python (version 1.3.1; Gramfort et al., 2013). To preserve the rank of the data, MaxFilter was not applied. Signal space projections were projected out for the sensor space analyses (Figure 2A-D) due to the strong influence of gradients on magnetoencephalographic recordings.

**Figure 2:**
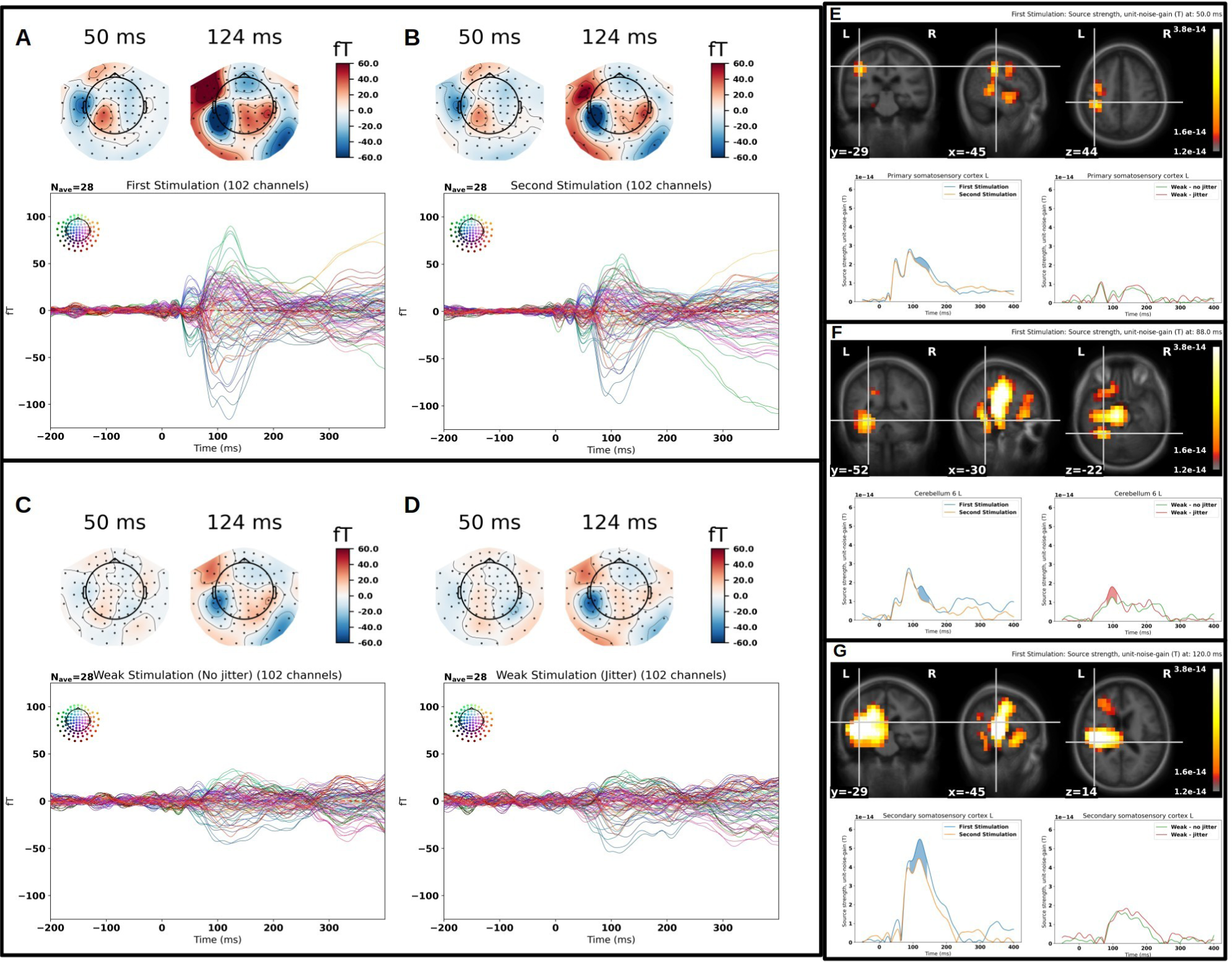
Evoked responses for the stimulation data and source localisation: The first and second stimulations are shown (**A&B**) revealing topographies compatible with primary (50 ms) and secondary (124 ms) somatosensory cortical activation. The weak stimulations are also shown (**C&D**), non-jittered and jittered. Only the secondary somatosensory cortical activation is apparent and weaker than for the first and second stimulations. For all topographies related to secondary somatosensory cortical activation (124 ms), posterior contributions are also seen. Bad channels have been interpolated. **E:** Primary somatosensory activation; for the first and second stimulations, we find a primary somatosensory activation at 50 ms. We find a difference cluster around the last peak (blue fill). The first response for the weak stimulations is at 60 ms. **F:** Cerebellar activation; for the first and second stimulations, we find three cerebellar peaks – at 49 ms, 87 ms and 124 ms respectively. We find a difference cluster around the last peak (blue fill). For the weak stimulations, we only find one peak, 97ms with a difference cluster (red fill). **G:** Secondary somatosensory activation; for the first and second stimulations, we find three peaks – at 52 ms, 88 ms and 120 ms respectively. We find a cluster around the last peak (blue fill). For the weak stimulations, we find an activation from 100 ms to 200 ms with no clear peaks. The vertices are chosen by finding the averaging the maximally responding vertices for First Stimulation and Second Stimulation and for Weak Stimulation (non-jittered) and Weak Stimulation (jittered) respectively within the relevant region of interest.

For the analysis of evoked responses, we low-pass filtered the data 40 Hz (one-pass, non-causal, finite impulse response; zero-phase; upper transition bandwidth: 10.00 Hz (-6 dB cutoff frequency: 45.00 Hz; filter length 331 samples; a 0.0194 passband ripple and 53 dB stopband attenuation). We subsequently segmented the filtered data into epochs of 1,200 ms, 200 ms pre-stimulus and 1,000 ms post-stimulus. The epochs were demeaned using the mean value of the pre-stimulus period. Bad channels were removed from each participant’s recording based on inspection of data. 4 participants had one magnetometer removed, and 6 participants had two magnetometers removed. For the sensor space analysis, segments of data including magnetometer responses greater than 4 pT (peak-to-peak) or gradiometer responses greater than 400 pT/cm (peak-to-peak) were rejected.

Subsequently epochs were averaged to create evoked responses for each of the trial types. On average, 291 (SD = 21.2) *First Stimulation*, 293 (SD = 18.2) *Second Stimulation*, 74 (SD =) *Weak, non-jittered*; 74 (SD = 3.35) *Weak, jittered*; 74 (SD = 2.20) *Omission, non-jittered;* 74 (SD = 3.93) *Omission, jittered* (SD = 3.53) remained. For the source space analysis, where signal space projections were not projected out, we did not apply this automatic rejection procedure, as the strong influence of gradients would mean that they would all be rejected out. This procedure is in line with recent advice for how to analyse electrophysiological data (Delorme, 2023).

For the analysis of oscillatory, beta band, responses, the data were band-pass filtered (14-30 Hz) (lower transition bandwidth: 3.50 Hz (-6 dB cutoff frequency: 12.25 Hz); upper transition bandwidth: 7.50 Hz (-6 dB cutoff frequency: 33.75 Hz); filter length 943 samples; passband ripple: 0.0194; stopband attenuation: 53 dB) bands. This filter was one-pass, non-causal, finite impulse response and zero-phase. We subsequently Hilbert-transformed the filtered data and segmented it into epochs of 1,500 ms, 750 ms pre-stimulus and 750 ms post-stimulus.

#### 2.5.1 Source reconstruction

A linearly constrained minimum variance beamformer was used to reconstruct sources for both the evoked and the oscillatory responses. For the evoked responses, the data covariance matrix was estimated based on the post-stimulus time period (from 0 ms to 1000 ms) and for the oscillatory responses the data covariance matrix was based on the whole period (from -750 ms to 750 ms). The sources for the oscillatory were estimated by taking the mean of the estimated source time courses for each segment for each band. Only magnetometers were used as these are the most sensitive to deeper sources. No regularisation was applied to the covariance matrices as they were well-conditioned. For estimating the spatial filter, we used the unit-noise-gain constraint, as this results in the best specificity for estimating the source time courses (Sekihara and Nagarajan, 2008), choosing the source orientation that maximises the signal-to-noise ratio. For both evoked and oscillatory responses, we took the absolute value of the source time courses.

To create a source space where the source time courses would be estimated from, we acquired sagittal T1-weighted 3D images for each participant using a Siemens Magnetom Prisma 3T MRI. The pulse sequence parameters were: 1 mm isotropic resolution; field of view: 256 x 240 mm; 192 slices; slice thickness: 1 mm; bandwidth per pixel: 290 Hz/pixel; flip angle: 9°; inversion time (TI), 1,100 ms; echo time (TE): 2.61 ms; repetition time (TR): 2,300 ms. We furthermore acquired T2-weighted 3D images. The pulse sequence parameters were: 1 mm isotropic resolution; field of view 256 x 240 mm; 96 slices; slice thickness: 1.1 mm; bandwidth per pixel: 780 Hz/pixel; flip angle: 111°; echo time: 85 ms; repetition time: 14,630 ms. Based on the T1 images a 1-layered boundary element method volume conductor was modelled based on delineating the inner skull using the watershed algorithm, as implemented in FreeSurfer. Based on the segmentation of the brain a volumetric source space was created with sources 7.5 mm apart. After the estimation of the source time courses in the individual source space, the source time courses were morphed onto *fsaverage*, a template from FreeSurfer (Dale et al., 1999; Fischl et al., 1999). All regions of interest were defined using the automated anatomical labeling atlas (AAL) (Tzourio-Mazoyer et al., 2002) except for secondary somatosensory cortex, which was defined using the Harvard Oxford cortical atlas (Desikan et al., 2006).

### 2.6 Statistical analysis

#### 2.6.1 Behavioural data

For the statistical analysis of the behavioural data, mixed effects models (Gelman and Hill, 2006) were fitted. We fitted logistic regression models with *Accuracy* as a binary dependent variable. For the final stimulation in a sequence, accuracy was coded as 1 if the participant indicated that they were stimulated, when they were in fact stimulated or if the participant indicated that they were not stimulated when they were in fact not stimulated. Accuracy was coded as 0 if the participant indicated that they were stimulated, when they were in fact not stimulated or if the participant indicated that they were not stimulated when they were in fact stimulated.

We had two independent variables of interest, *Stimulation Type* and *Stimulation Variance*. Stimulation Type was a factorial variable with two levels, *Omission* or *Weak*. Stimulation Variance was a continuous variable expressing the variance around the expected time of stimulation for the jittered stimulations, i.e. the fourth, fifth and sixth stimulations. The Stimulation Variance was calculated as:

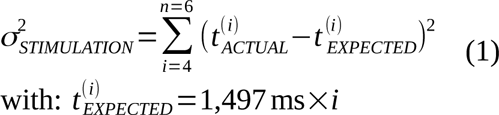

*t_ACTUAL_* was when the stimulation actually happened. This resulted in a Stimulation Variance equal to zero for the non-jittered sequences and a variable Stimulation Variance for the jittered sequences.

For the subject-level effects (random effects), we modelled a unique intercept for each subject and two unique slopes of the Stimulation Variance as dependent on the Stimulation Type, Omission or Weak. For the group-level effects (fixed effects), we followed the strategy of beginning with a null model that only estimated a mean accuracy (an intercept). Subsequently, we added the independent variables of interest in a step-wise fashion to the model – for each step calculating the likelihood-ratio between the two models:

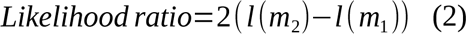

with *l* as the log-likelihood and *m* as the models (*m_1_* is nested with *m_2_*). This ratio is approximated by a *χ*²-distribution. Whether the null hypothesis, that the likelihood ratio is 0 could be rejected, was evaluated using a *χ*²-test with the likelihood ratio as the test statistic and degrees of freedom equal to the difference in free parameters between the models. We rejected the null hypothesis at *α* = 0.05.

The mixed models were estimated using the *lme4* package (Bates et al., 2014) from *R* (R Core Team, 2020).

#### 2.6.2 Magnetoencephalographic data

For the statistical analysis of the magnetoencephalographic data, we focused on the regions and time intervals reported by Andersen and Dalal (2021), i.e. the cerebellum, and the primary and secondary somatosensory areas. We used a linearly constrained minimum-variance beamformer (Sekihara and Nagarajan, 2008) to extract time courses from each of these areas.

For evoked responses, the trial type comparisons of interest were between *First Stimulation* and *Second Stimulation*, between *Weak, no jitter* and *Weak, jittered*, and between *Omission, no jitter* and *Omission, jittered* in the time interval between 0 ms and 150 ms. Two time courses were extracted, namely the ones that showed the maximal response for each condition within the region of interest. We ran cluster permutation statistics on the mean of the extracted time courses with 10,000 permutations (Maris and Oostenveld, 2007), using *α* = 0.05.

For the beta band (14-30 Hz) oscillatory responses of the omissions, the trial type comparisons of interest were the same. The time interval run on was 0-200 ms based on the findings of Andersen and Dalal (2022, 2021). In a separate analysis in the beta band, we investigated the contributions of thalamus as a potential contributor to stimulus processing, as this was also found by Andersen and Dalal (2022, 2021).

#### 2.6.3 Correlations between behavioural and magnetoencephalographic data

To estimate correlations between brain activity and behaviour, we, based on the winning logistic regression model, extracted the variance-related slope for each participant for each task condition, i.e. for *Weak* and *Omission*. These were then to be correlated with relevant brain activity. The more negative the slope, the more a given participant’s performance would deteriorate as a function of the variance introduced in the jittered conditions. Spearman correlations were run as the data was non-normal.

#### 2.6.4 Envelope correlations

To investigate oscillation-mediated connectivity between regions of interest, we used envelope correlation (Hipp et al., 2012) that as its name indicates estimates the correlation of envelopes of oscillatory activity. We defined the cerebellum as a seed and investigated its connections to the the rest of the brain and the thalamus in particular. We conducted an analysis of variance using two factors, Stimulation Type (2 levels: weak, omission) and Regularity (2 levels: non-jittered, jittered). We investigated the envelope correlations from -100 ms to 100 ms around *Weak* and *Omission* in the beta band. This was done as an exploratory analysis.

## 3 Results

### 3.1 Behaviour

Adding Stimulation Type as a group-level variable to the null model, which only modelled mean accuracy besides the subject-level effects, resulted in a significant increase in the likelihood of the model, *χ*²(1) = 13.8, *p* = 0.0020. Further adding Stimulation Variance to this model also resulted in a significant increase in the likelihood of the model, *χ*²(1) = 5.92, *p* = 0.014. Adding the interaction between Stimulation Type and Stimulation Variance did, however, not result in a significant increase in the likelihood of the model, *χ*²(1) = 0.0083 *p* = 0.93. Our winning model thus includes group-level effects of Stimulation Variance and Stimulation Type, but no interaction between them. Given this model, the estimated group-level accuracy when Stimulation Variance was 0 (no jitter) for Omission was 91.7 %, and for Weak it was 78.7 % (Figure 3A). At the mean Stimulation Variance of the jittered trials (0.032 s²), the estimated group-level accuracy had dropped to 90.7 % for Omission and to 76.5 % for Weak. At the maximum Stimulation Variance (0.09 s²), the estimated group-level accuracy had dropped to 88.4 % for Omission and to 71.6 % for Weak (Figure 3A). We can thus conclude that our paradigm had the intended effect on behaviour, resulting in detection of stimuli becoming worse, the more jitter there was in the stimulations leading up to the target stimulation.

**Figure 3:**
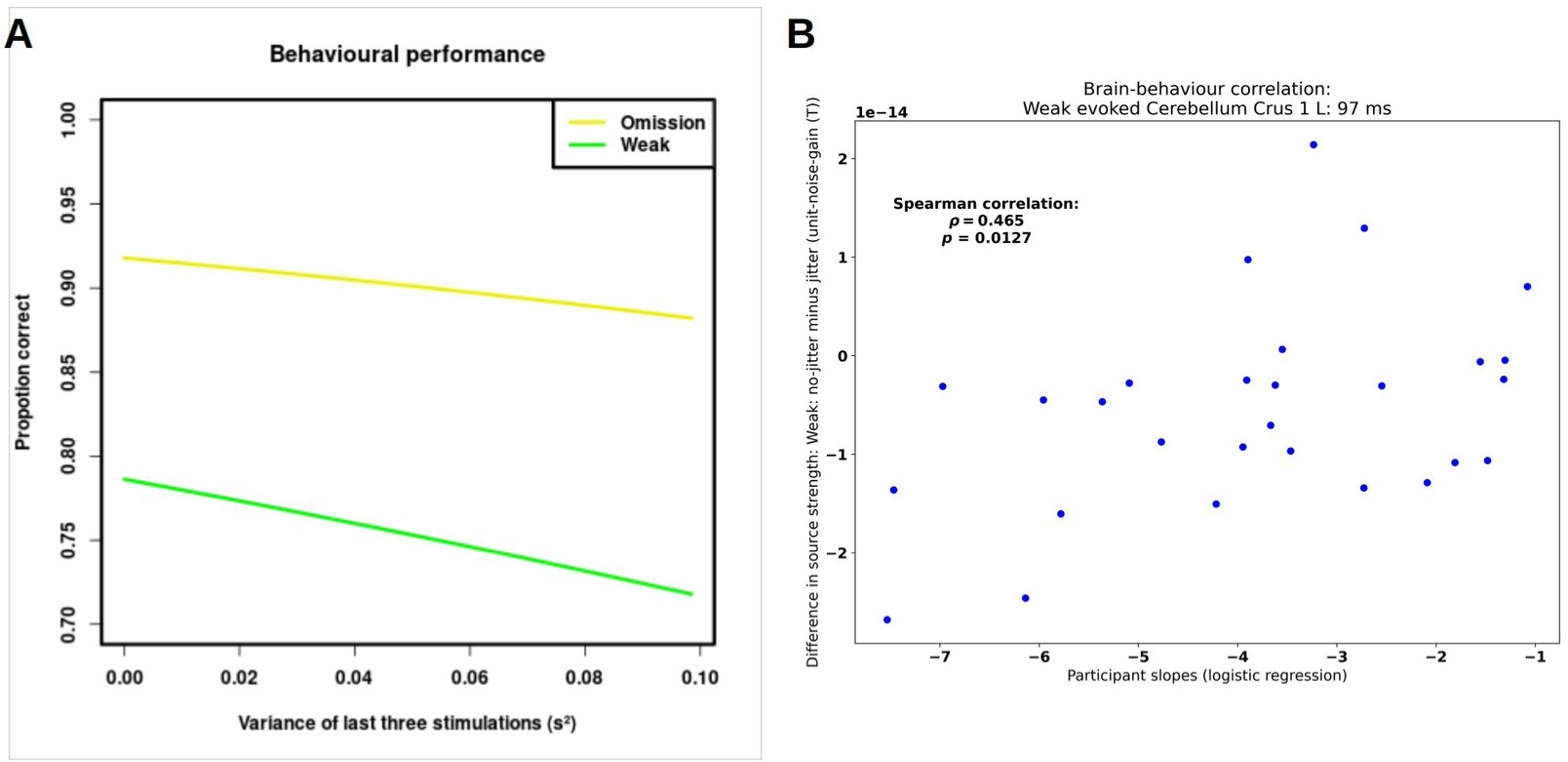
Behavioural performance and correlations between the estimated participant slopes based on multilevel logistic regression and the difference in estimated peaks for the cerebellum for the Weak Stimulations (no-jitter minus jitter) and the Omitted Stimulations (no-jitter minus jitter): **A:** proportion correct as dependent on the variance of the last three stimulations. **B:** we find a significant positive correlation between the estimated participant slopes and the difference between the evoked responses to the Weak Stimulations.

### 3.2 Magnetoencephalography

#### 3.2.1 Evoked responses

For the evoked responses, we found the expected responses compatible with primary and secondary somatosensory cortical activations (Figure 2). The topographies related to the secondary somatosensory activation also included posterior contributions, potentially compatible with cerebellar contributions. The linearly constrained minimum variance beamformer revealed the expected primary and secondary somatosensory cortical activations for First Stimulation. Furthermore, activations in left cerebellar lobule VI and left cerebellar crus I were also revealed (Figure 2F). We used the coordinates associated with the maximum activation for each of these regions of interest to estimate the responses associated with Weak, non-jittered and Weak, jittered. Running the cluster permutation statistics revealed six significant effects of condition. The first four ones were for the comparison between First Stimulation and Second Stimulation in the primary somatosensory cortex, *p* = 0.0004, secondary somatosensory cortex, *p* = 0.0001 and in the left cerebellar lobule VI *p* = 0.0088, and left cerebellar crus I, *p* = 0.0084. The clusters associated with this extended from 106 ms to 149 ms, from 90 ms to 142 ms, from 116 ms to 149 ms, and from 116 ms to 149 ms. respectively. The two last ones were for the comparisons between Weak, non-jittered and Weak, jittered in the left cerebellar lobule VI, *p* = 0.0124 and the left cerebellar crus I, *p* = 0.0295. The clusters associated with this extended from 85 ms to 113 ms and from 87 ms to 111 ms, respectively.

##### 3.2.1.1 Brain-behaviour correlations – evoked responses

We correlated the participant level slopes for the *Weak Stimulation* condition, estimated from the winning behavioural model, with the differences (between *Weak, no jitter* and *Weak, jittered*) in peak evoked cerebellar crus I activity (97 ms) (Figure 2G), *ρ*_26_ = 0.465, *p* = 0.0127 (Figure 3B). (cerebellar lobule VI showed a similar effect, *ρ*_26_ = 0.374, *p* = 0.0497). The positive sign of *ρ* indicates that the more positive the difference in cerebellar source strength between Weak, non-jittered and Weak, jittered, the less negative were the participant slopes related to variance in the jittered condition. In other words, stronger cerebellar activation to a jittered weak stimulus than to a non-jittered weak stimulus was correlated with *worse* performance.

#### 3.2.2 Oscillatory responses

##### 3.2.2.1 Involvement of the beta band (14-30 Hz) in prediction

Running the cluster permutation statistics, we reject the null-hypothesis that the non-jittered omissions and jittered omissions are exchangeable, *p =* 0.0127. We replicate the finding from Andersen and Dalal (2022, 2021), as we find a cluster (Figure 4) that contains left cerebellar lobule VI extended from 30 ms to 65 ms and from 98 ms to 115 ms and left cerebellar crus I, extended from 29 ms to 52 ms. This also includes left primary somatosensory cortex, (176 ms to 200 ms), and left secondary somatosensory cortex, (154 ms to 179 ms).

**Figure 4:**
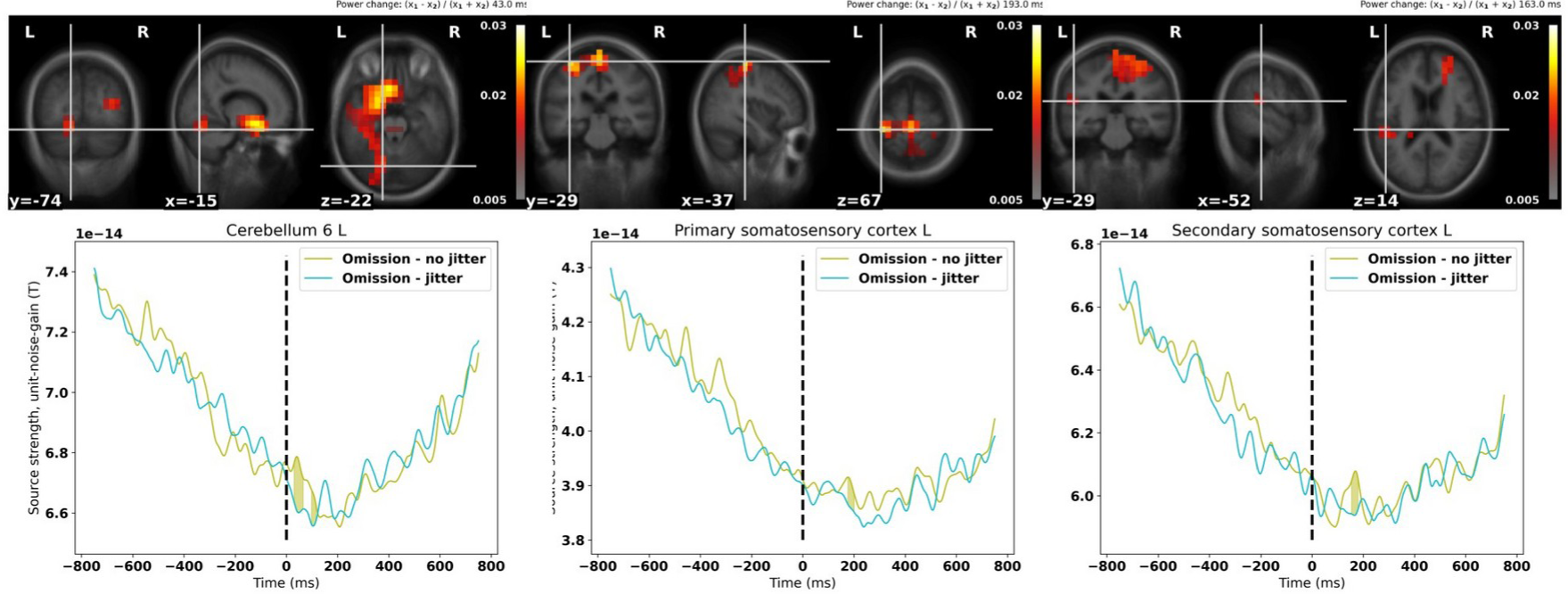
Replication of results from Andersen and Dalal (2022, 2021); *cerebellar lobule VI shows increased activity just after the time of the expected stimulation for non-jittered omissions compared to jittered omissions. Furthermore, primary and secondary somatosensory cortices are also involved with secondary somatosensory cortex showing involvement after that of cerebellar lobule VI and primary somatosensory cortex. The time courses are extracted from the voxel shown in the brain maps and the solid filling show the temporal extent of the cluster. In the brain maps, only voxels that part of a cluster are shown, all other voxels are masked. Voxels show the power change between non-jittered Omissions and jittered Omissions (x1 – x2) / (x1 + x2)*.

For the comparison between First Stimulation and Second Stimulation, we reject the null-hypothesis that they are exchangeable, *p* = 0.000977. Peaks are found in the primary and secondary somatosensory cortices and the thalamus (Figure 5). These are separated in space supporting their veracity. The thalamic peak is explored further in section 3.4 below.

**Figure 5:**
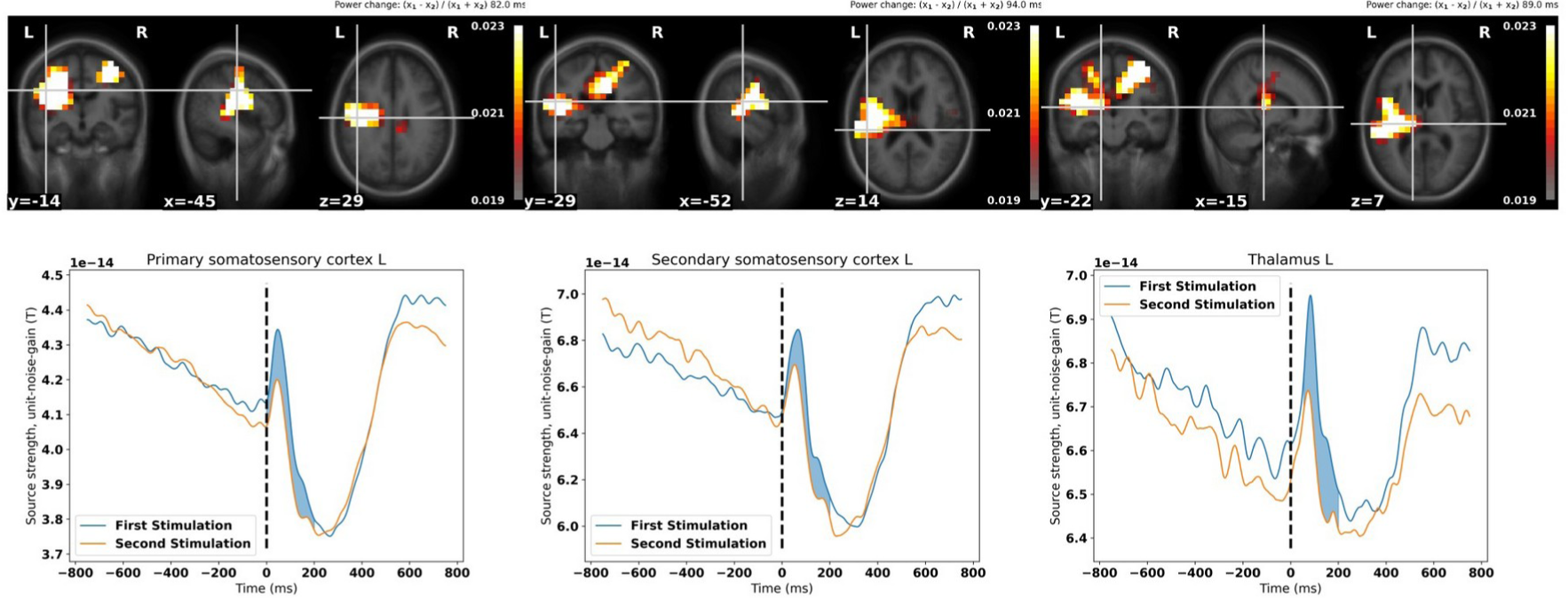
Power change between First Stimulation and Second Stimulation in the beta band (14-30 Hz): *we find peaks in the left primary, 47 ms, and secondary, 67 ms, somatosensory cortices, and in the thalamus, 84 ms. The time courses are extracted from the voxel shown in the brain maps and the solid filling show the temporal extent of the cluster. In the brain maps, only voxels part of a cluster are shown, all other voxels are masked. Voxels show the power change between First Stimulation and Second Stimulation (x1 – x2) / (x1 + x2)*.

### 3.3 Envelope correlations

The envelope correlations in the beta band revealed differences in correlations for jittered and non-jittered weak stimulations and omitted stimulations between left cerebellar crus I and the right thalamus. An analysis of variance revealed an interaction between Stimulation Type (2 levels) and Regularity (2 levels), *F*_1,100_ = 4.42, *p* = 0.0376 for the right thalamus (Figure 6ABC).

**Figure 6:**
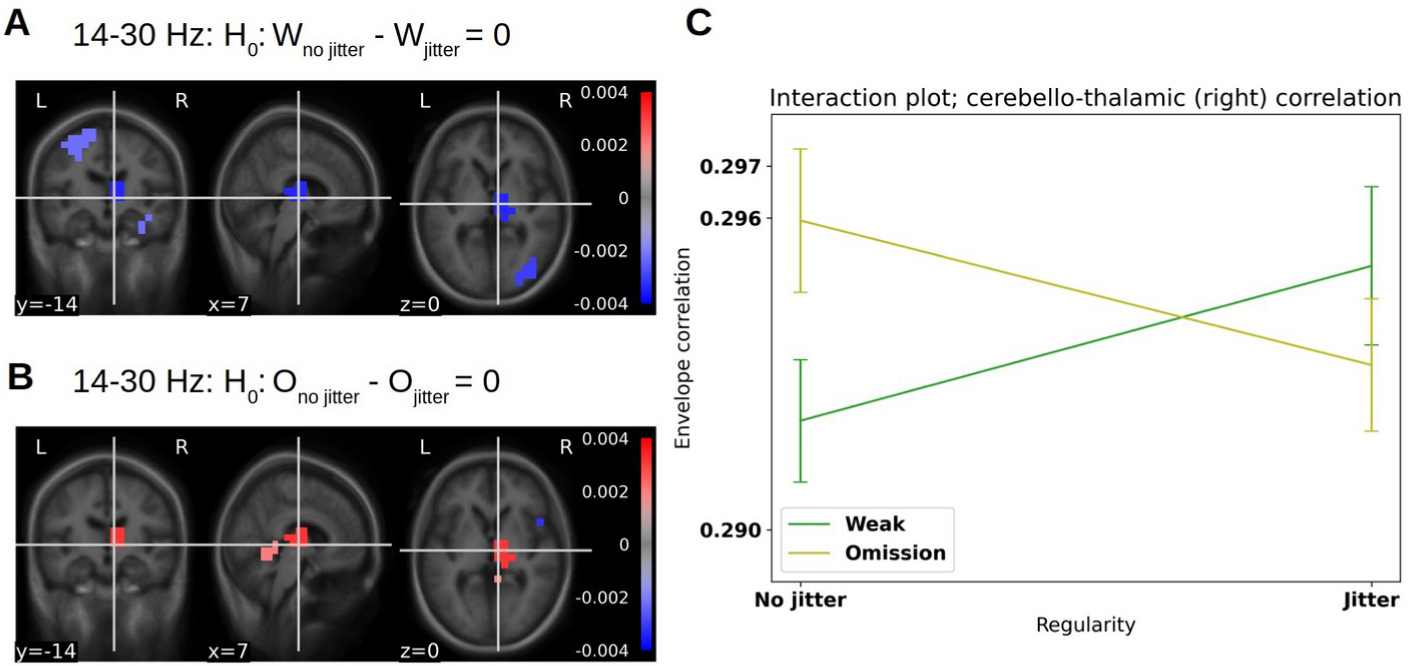
Cerebello-thalamic envelope correlations: Envelope correlation using left cerebellar crus I as a seed: only envelope correlations in the beta band (14-30 Hz) that are significant at α = 0.05 are shown. **A:** Envelope correlations are stronger to the thalamus for the non-jittered condition for the weak stimulations, whereas **B:** envelope correlations are stronger to the thalamus for the jittered condition for the omissions (bottom right). **C:** interaction plot for the right thalamus, highlighting the increase in correlation for weak stimulations when introducing jitter and the decrease in correlation for omitted stimulations when introducing jitter. Error bars are standard errors of the mean.

Beta band envelopes in cerebellar crus I correlated differently with the right thalamus dependent on whether the last stimulus was omitted or weak, while this in turn depended on whether or not the sequence was jittered. Specifically, we found that cerebello-thalamic envelope correlations *increased* for Weak Stimulations when going from non-jittered to jittered, whereas these *decreased for* Omitted Stimulations when going from non-jittered to jittered (Figure 6C). See supplementary material 1 for other significant modulations.

### 3.4 Is it really the thalamus?

To corroborate that the thalamic findings are not due to the spread of activity from other areas, we investigated the evoked responses and the source reconstructions more closely (Figure 7).

**Figure 7:**
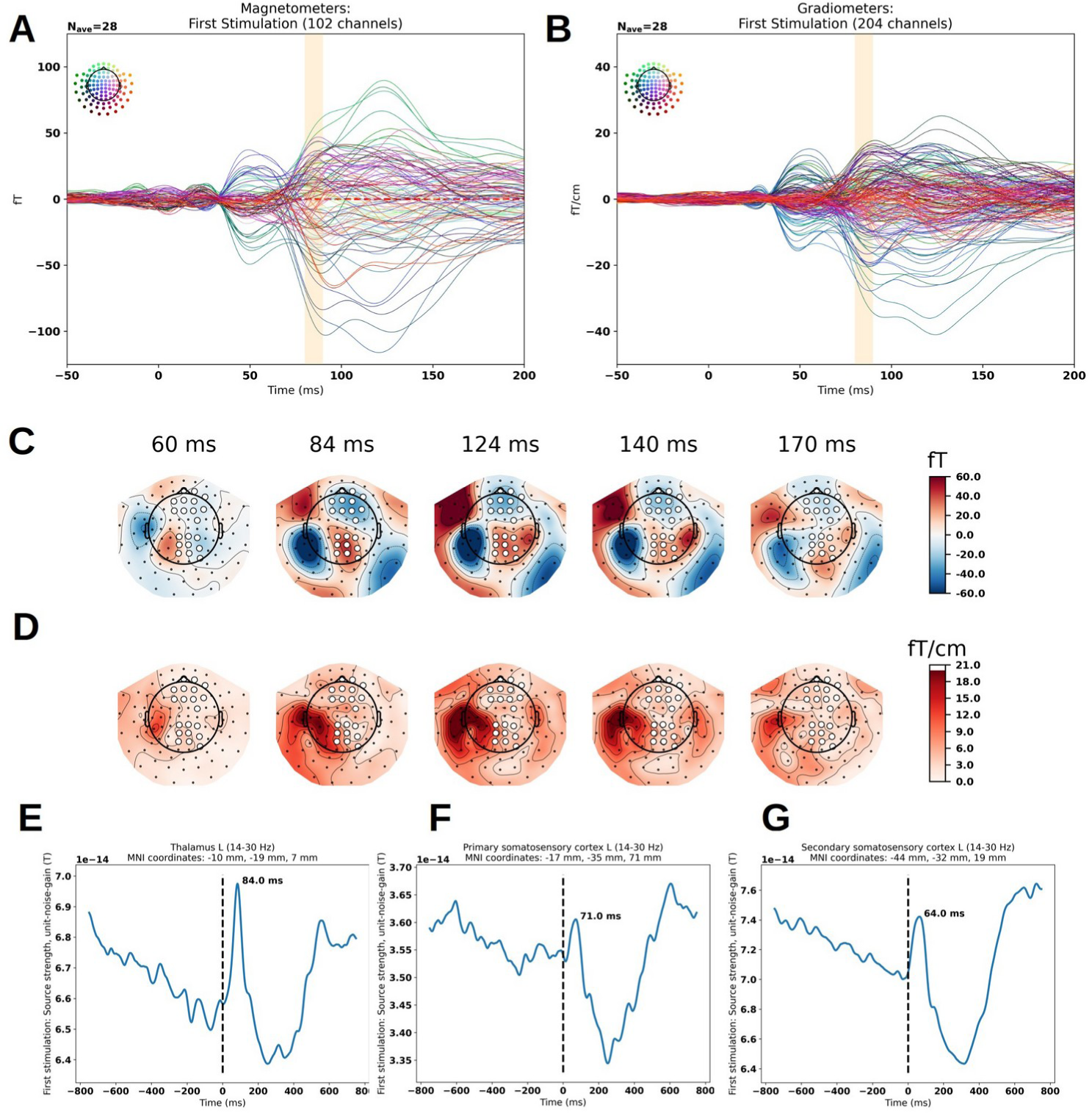
Compatibility of the grand averaged evoked response with the beta band (14-30 Hz) thalamic response found. **A:** magnetometers: a component is highlighted at 84 ms for the first stimulations for the evoked responses (not discussed in Figure 2). **B:** planar gradiometers: the same component is highlighted at 84 ms. **C:** the topographical plots for the magnetometers are consistent with two underlying sources at 84 ms, one lateral and one central (highlighted in white); at 124 ms they are still both present, whereas at 140 ms the lateral one starts dominating and at 170 ms, it is the only one that remains. **D:** on the topographical plots for the gradiometers, the central component is absent, consistent with a deep central source in **C** (same sensor triplets highlighted in white) **E:** the peak of the beta band response localised to the left thalamus match the peaks in the evoked response (**A**&**B**). **F:** the beta band response of the primary somatosensory cortex peaking earlier, 71 ms, than the thalamic response. **G:** the beta band response of the secondary somatosensory cortex peaking earlier, 64 ms, than the thalamic response. The Montreal Neurological Institute (MNI) coordinates for these estimated sources were chosen based on finding central coordinates for the relevant regions from the MNI brain from FSL (Smith et al., 2004).

## 4 Discussion

In summary, we found that the jitter introduced in a sequence affected the detection of weak stimulation (Figure 3A). Furthermore, we found cerebellar responses evoked by the stimulations (Figure 2). For the weak stimulations, the difference in estimated source current density between jittered and non-jittered conditions correlated with the scale of the slope indicating how much participant performance was affected by variation in the stimulation sequences (Figure 3B). In the beta band, we furthermore found differences in the primary and secondary somatosensory cortices and the thalamus between the first, surprising, stimulation and the second, expected, stimulation (Figure 5). Finally, using the cerebellum as a seed we found that the envelope of cerebellar crus I correlated with the envelope of right thalamic activity (Figure 6AB).

The finding that the variance introduced in the behavioural task in the jittered condition makes it harder to detect the weak stimulations, which are otherwise identical, corroborates that there must be an internal mechanism that tracks the expected timing of stimulation (Figure 3A). The replication of the beta band finding that cerebellum clocks the timing of expected stimulation corroborates the finding of Andersen and Dalal (2022, 2021) and also highlights the fact that it can be done with half the trials (75 instead of 150) (Figure 4). This establishes beta band cerebellar lobule VI activity as following expected patterns of stimulation and establishes cerebellum as a candidate for enabling the better performance seen when stimulation sequences are regular.

That the proposed cerebellar clocking activity has functional relevance is evidenced by the correlation (Figure 3B) found between differences in cerebellar evoked responses (Figure 2) for weak stimulations depending on whether they are preceded by a jittered sequence or not. Interestingly, the strength of the cerebellar evoked response does not enable stronger performance on the behavioural task, but the opposite: the stronger the cerebellar response to the jittered weak stimulation was relative to the non-jittered one, the worse subject performance was as estimated by the slope of their individual fit. This is evidence that the cerebellum is not merely encoding the perceived magnitude of the electrical stimulation – on the other hand, the increased cerebellar response for the jittered sequences may indicate inhibition of weak stimulation following a jittered sequence. The less inhibited a weak stimulation was, i.e. the smaller the difference between weak cerebellar evoked responses (Figure 4B; *y*-axis), the greater the probability was of detecting it (Figure 4A). In fact, the Purkinje cells, which are the neurophysiological basis of magnetoencephalographic signals (Samuelsson et al., 2020), have an inhibitory role in sensory processing and perception (D’Angelo, 2018; D’Angelo et al., 2009; Ito et al., 1970). The role of this particular mechanism could be to filter away non-salient crossings of the sensory threshold, signalling whether a stimulation took place or not. According to this interpretation, weak stimulation following a jittered sequence would have a higher chance of being filtered away, thereby leading to more *misses* as variance increases (Figure 3A). This interpretation of cerebellar activity as inhibitory is also consistent with the impact of jitter on performance for omissions. That beta band power is lower for omissions following a jittered sequence implies that any non-salient crossings of the threshold are less likely to be filtered away. As a consequence, more *false alarms* should follow as variance increases, which is indeed what we observe (Figure 3A).

We also found a difference in beta band activity between first and second stimulations – the thalamic responses are similar to the ones reported by Andersen and Dalal (2022: Supplementary Material 3: New Figure 3). This justified running the exploratory functional connectivity analysis of the cerebellum and the thalamus, where we found an interaction in the envelope correlations between left cerebellar crus I and the right thalamus dependent on stimulation type and regularity of the preceding sequence. Assuming that this connection reflects the degree to which the cerebellar expectations are relevant to informed behaviour, the overall decrease in connectivity for jittered weak stimulation compared to non-jittered weak stimulation may explain the decrease in performance in detecting weak stimulation when jitter is introduced (Figure 4B). Put in terms of signal-detection theory, when the number of *hits* decreases and thus the number of *misses* increases, the connectivity increases. And vice versa, when the number of *correct rejections* decreases, the number of *false alarms* increases, the connectivity decreases. That means that a decrease in connectivity between left cerebellar crus I and the right thalamus corresponds with answering stimulus-present and an increase in cerebello-thalamic connectivity answering no-stimulus-present, independently of whether a stimulus was actually present. This can be interpreted as cerebellum connecting more strongly with thalamus when there is no perceived stimulus, in line with the idea that the cerebellum detects omissions from the established pattern, which results in increased beta band power (Figure 4). This interpretation of the involvement of right thalamus in motor commands is consistent with the contralaterality of thalamic efferents (Herrero et al., 2002), i.e. subjects indicated their answers with their left hand. The timing of the response may seem late, as the first-order thalamic relay-activity happens early, < 20 ms. However, it has been argued that higher-order relay-activity is likely to happen at later stages in the processing of stimuli, especially if they are relevant to motor output (Guillery and Sherman, 2002; Sherman, 2007).

As appealing as these thalamic and cerebello-thalamic results are, it is important to treat magnetoencephalographic results of thalamus with a critical attitude. The thalamus is a non-optimal target for magnetoencephalography due to its deep location towards the centre of the head and the configuration of its neurons, which do not form a field as open as those of the cerebral and cerebellar cortices (Lorente de Nó, 1947), even though simulation and experimental studies have also been used to argue for the feasibility of detecting the thalamus using magnetoencephalography (Attal et al., 2012, 2009; Attal and Schwartz, 2013). Therefore, we scrutinised the thalamic responses more carefully (Figure 7) to make sure that the results were not just due to a spread of activity from nearby regions as a result of our source localisation procedure. Several points are indicative of this not being the case. Firstly, the thalamic peak response at 84 ms for the beta band (14-30 Hz) (Figure 7E) coincides with an evoked response at 84 ms visible in sensor space (Figure 7AB). Secondly, the topographical pattern at 84 ms for the magnetometers is compatible with a central, deep source (Figure 7C) (and a lateral source, the secondary somatosensory cortex). Thirdly, the topographical pattern at 84 ms (or later) for the planar gradiometers does not include a similar, central activation, but only a lateral one (Figure 7D). Fourthly, neither the primary somatosensory cortex nor the secondary somatosensory cortex peaked at a similar time for the beta band (Figure 7FG). We do not mean to imply that this means that the 84 ms peak of thalamic beta band activity is reflective of underlying oscillations – it may be that the focusing on the beta band makes it easier to separate the potential thalamic response from the concurrently unfolding secondary somatosensory cortex evoked response (Figure 7ABCD), which is represented at a lower frequency.

In conclusion, our study shows that evoked cerebellar responses may have an inhibitory role (Figure 2). These cerebellar responses have functional relevance, as they correlate with performance in detecting weak stimuli (Figure 3). Furthermore, cerebellar beta band (14-30 Hz) responses encode predictions (Figure 4), and connectivity results suggest that the evaluation of these cerebellar predictions inform behaviour through connectivity to the thalamus (Figure 6).

These findings highlight the need to include non-cortical areas (Andersen et al., 2020; Shine et al., 2023) in our understanding of the brain and its capabilities in prediction and using the environment to inform behaviour.

## 5#Acknowledgements

We thank Ida Bang Hansen for assistance in data collection. Lau M. Andersen was funded by the Lundbeck Foundation (R322-2019-1841). A portion of this project was funded by a European Research Council Starting Grant (640448) to Sarang S. Dalal.

