## Supplementary material for "Detection of threshold-level stimuli modulated by temporal predictions of the cerebellum": Supplematary tables 1 and 2

***Supplementary table 1:*** *areas from the  with functional connectivity to left cerebellar crus I, significant at α=0.05 for the difference between Weak, non-jittered and Weak, Jittered*

| Region of interest | t-value (df=26) | p-value |
|---|---|---|
| Precentral_L | -2.09 | 0.046 |
| Occipital_Mid_R | -2.33 | 0.027 |
| Occipital_Inf_R | -2.45 | 0.021 |
| Fusiform_R | -2.75 | 0.010 |
| Thalamus_R | -2.21 | 0.036 |

***Supplementary table 2:*** *areas from the  with functional connectivity to left cerebellar crus I, significant at α=0.05 for the difference between Omission, non-jittered and Omission, Jittered*

| Region of interest | t-value (df=26) | p-value |
|---|---|---|
| Frontal_Inf_Oper_L_L | -2.36 | 0.025 |
| Frontal_Inf_Oper_R | -2.41 | 0.023 |
| Occipital_Sup_R | -3.07 | 0.005 |
| Thalamus_R | 2.29 | 0.03 |
| Cerebellum_4_5_R | 2.15 | 0.04 |
